# Tumour rewires host tryptophan metabolism to promote organism-wide immune suppression

**DOI:** 10.64898/2026.09.02.748262

**Authors:** Lucia Garcia-Lopez, Ernesto Saez-Carrion, Mary L. Uribe, Carmen Gomez-Escolar, Esther Ballesta-Illan, Jason M. Tennesen, Maria Dominguez

**Affiliations:** Instituto de Neurociencias CSIC-UMH, Alicante, Spain; Indiana University, Department of Biology, Bloomington, IN 47405, USA; Member, Melvin and Bren Simon Cancer Center, Indianapolis, Indiana 46202, USA

**Keywords:** kynurenine pathway, 3-hydroxykynurenine, tumour–host communication, immunometabolism, immune tolerance, PTEN

## Abstract

How tumour genotype reshapes systemic metabolism to drive immune tolerance and tumour progression remains unclear. Here, we show that *Pten*-deficient *Drosophila* tumours reprogram host tryptophan metabolism, triggering systemic and local changes in immune compartments to foster tolerance. Tumour-derived nitric oxide activates the kynurenine pathway in liver-like tissue, selectively increasing the 3-hydroxykynurenine (3-HK) branch. Host 3-HK promotes tumour progression and suppresses immune function via aryl hydrocarbon receptor (AhR) signalling. These results identify the kynurenine pathway as a central tumour–host nexus and reveal how tumour genotype creates immune vulnerabilities.

## Main

Cancer is a systemic disease that profoundly remodels host physiology. Local tumour progression is shaped not only by the tumour’s genotype but also by dynamic interactions with host metabolism^1^, inflammation^2^, immunity^3,4^, and nutrient status^5,6^. Tumours and host tissues compete for resources, triggering widespread metabolic and physiological reprogramming well beyond the primary site. However, whether specific oncogenic mutations direct unique systemic immunometabolic states, or if these are simply consequences of tumour burden, remains unresolved.

Although innate immunity efficiently eliminates damaged and pre-malignant cells^7,8^, the host ultimately faces a trade-off: persistent tumour-associated inflammation, if left unchecked, results in excessive oxidative stress, collateral tissue damage, and metabolic disruption^9^. To limit these detrimental effects, organisms engage immune tolerance programmes that restrain excessive immune activation, even at the expense of permitting abnormal cell survival^10^. Tumours can exploit these physiological tolerance mechanisms to evade immune destruction. Although current immunotherapies can restore anti-tumour immunity, their efficacy varies widely across patients and tumour types and is frequently accompanied by autoimmune-related toxicity^11^. These variable outcomes indicate that immune evasion cannot be explained by a single suppressive mechanism. Rather, the remarkable diversity of tumour immune microenvironments and the multitude of immunosuppressive pathways that operate within them are likely to determine both tumour progression and responses to immunotherapy^12^.

A major regulator of immune tolerance is the evolutionarily conserved kynurenine pathway of tryptophan metabolism^13,14^, which acts largely through aryl hydrocarbon receptor (AhR) signalling^15^. In mammals, indoleamine 2,3-dioxygenases (IDOs), induced in many cancers, together with the liver-enriched enzyme tryptophan 2,3-dioxygenase (TDO), initiate tryptophan degradation to kynurenine. Downstream metabolites exert diverse immunomodulatory, cytoprotective and cytotoxic activities^14^. Although altered kynurenine metabolism has been extensively linked to immune suppression in advanced cancers and *in vitro*, how tumours reprogramme systemic tryptophan metabolism to coordinate host immunity and metabolism during *in vivo* tumour progression remains poorly understood.

*PTEN*-deficient cancers are characterised by profound metabolic rewiring^16^, immune evasion^17,18^, and *PTEN* loss is associated with poor responses to immunotherapy^19^. Intriguingly, in both flies and mice, dietary restriction paradoxically accelerates, rather than suppresses, *PTEN*-deficient tumour growth^6,20^, suggesting that tumour genotype can induce maladaptive systemic responses by the host. Understanding the mechanisms underlying this paradox may reveal how tumour–host metabolic interactions determine disease progression and the outcome of dietary or immunomodulatory interventions.

*Drosophila melanogaster* provides a powerful *in vivo* model to investigate tumour–host interactions because genetic, metabolic, and immune pathways governing cancer progression are conserved with mammals^5^. *Pten*-deficient tumours in flies recapitulate key hallmarks of mammalian cancer, including metabolic reprogramming, inflammation, and therapeutic resistance^6,18,21,22^. The fly innate immune system, comprising plasmatocytes, crystal cells and lamellocytes, performs functions analogous to mammalian innate immunity, including phagocytosis, cytotoxic responses, and inflammatory regulation^23,24^. Moreover, the evolutionary conservation of the tryptophan–kynurenine pathway (Fig. 1a) provides an opportunity to determine how tumour genotype remodels systemic metabolism and host immunity.

**Fig. 1.**
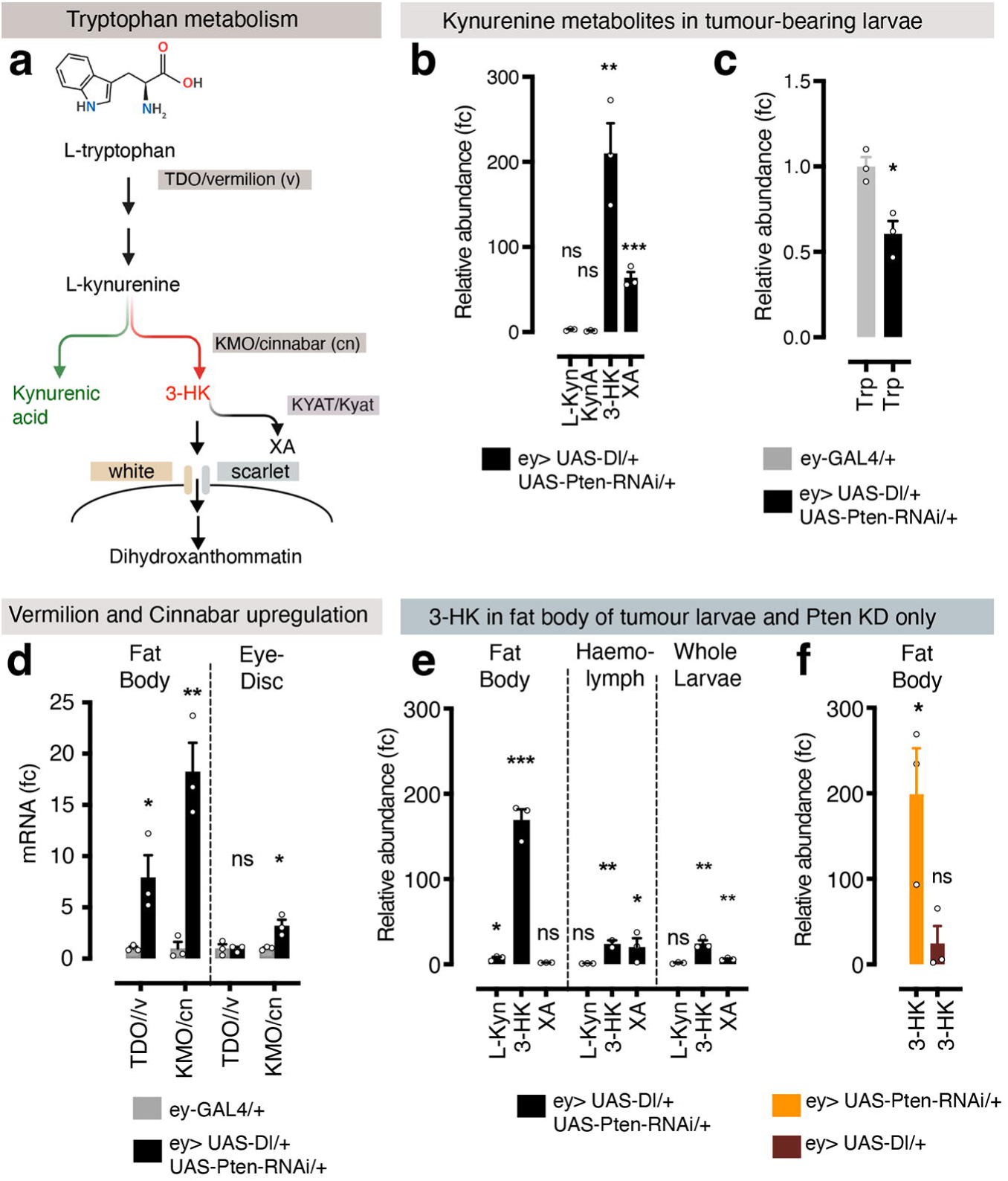
Systemic tryptophan metabolism rewiring in *Pten*-deficient tumours. **a,** Schematic of the conserved kynurenine pathway (KP) in mammals and *Drosophila*, showing the enzymatic conversion of tryptophan to kynurenine and its subsequent metabolism through the 3-hydroxykynurenine (3-HK) and kynurenic acid (KynA) branches. The *Drosophila* homologues vermilion (TDO) and cinnabar (KMO), predominantly expressed in the fat body, generate 3-HK, which is transported by the White–Scarlet ABC transporter to pigment cells for ommochrome synthesis^54^. **b,c,** Untargeted UHPLC–MS metabolomic profiling of third instar larvae (wandering stage) reveals a marked elevation of the tryptophan-derived metabolite 3-HK in tumour-bearing larvae (**b**) and a modest reduction of systemic tryptophan (**c**) relative to control larvae (*ey-GAL4*: *n* = 3 biological repeats and 15 larvae per sample). Multiple unpaired t-test, from left to right: *P =* 0.004229 (**), *P* = 0.000793 (***). For comparison of tryptophan levels between larvae *ey-GAL4* and *ey>Dl>Pten RNAi/+* an unpaired two-tailed Student t-test was performed: *P* = 0.0134 (*). In this and other figures, genotypes are indicated by the colour-coded keys below the graphs; genotype–colour assignments are consistent throughout the figures. **d,** qRT–PCR analysis of *TDO/vermilion* and *KMO/cinnabar* in dissected fat bodies or eye-antennal discs of tumour-bearing larvae compared to control *ey-GAL4* larvae. Bar plots indicate mean ± s.e.m. (*n* = 3), with individual data points shown. Expression levels are normalised to the *Rp49* housekeeping gene and presented as fold-change relative to *ey-GAL4* controls. Statistical comparisons were performed using unpaired two-tailed Student’s t-tests (left to right): *P* = 0. 0335 (*), *P* = 0.0039, *P* = 0.0202 (*). **e, f,** GC–MS analysis of tumour-bearing larvae (fold change to control *ey-GAL4*) of dissected fat bodies, haemolymph, or whole larvae. Bar plots show relative abundance (fold change to control *ey-GAL4/+*) and presented as mean ± s.e.m., with dots representing individual values (*n* = 2 or 3 biological repeats, see Source Data). Data was analysed using an unpaired two-sided Student t-test (from left to right): *P* = 0.0204 (*), *P* = 0.0002 (***), *P* = 0.0049 (**), *P =* 0.0053 (**), *P =*0.0188 (*), *P =* 0,0031 (**), and *P* = 0.0042 (**). **f,** Accumulation of 3-HK in the fat body of larvae with *Pten* knockdown via RNAi only or *Dl* overexpression only. Bar plots show relative abundance (fold change to control *ey-GAL4/+*) and presented as mean ± s.e.m., with dots representing individual values. Ordinary one-way ANOVA was performed to control (*ey-GAL4*) to *ey>Pten RNAi* and *ey>Dl* alone. *P* = 0.0133 (*).

We previously showed that *Drosophila Pten* loss cooperates with activated Notch to drive malignant tumour progression in both flies and human cancer and that constitutive PI3K–Akt signalling resulting from loss of *Pten* promotes inflammation accompanied by an apparent suppression of early anti-tumour immunity^18,21^. Here, we show that *Pten*-deficient tumours actively impose systemic immune tolerance by remotely reprogramming host tryptophan metabolism through tumour-derived nitric oxide (NO). This tumour–host signalling axis amplifies kynurenine pathway activity in the fat body, leading to accumulation of the pro-oxidant metabolite 3-hydroxykynurenine (3-HK), suppression of cytotoxic melanisation responses, and extensive remodelling of the immune microenvironment into a tolerogenic state that supports tumour growth while limiting host inflammatory damage. Together, our findings identify tumour-driven rewiring of host tryptophan metabolism as a mechanism by which tumour genotype orchestrates systemic metabolic and immune adaptation, uncovering a targetable pathway that links cancer metabolism with systemic immune tolerance.

## Results

### *Pten*-deficient tumours induce systemic activation of the 3-hydroxykynurenine branch of tryptophan metabolism

How localised oncogenic lesions reprogramme whole-organism physiology, and how these systemic responses influence tumour progression, remain fundamental questions in cancer biology. To address these questions, we used a *Drosophila* model with *Pten* knockdown and Notch activation in the developing eye epithelium^18^ (*ey>UAS-Delta (Dl)>UAS-Pten RNAi*; hereafter referred to as oncogenic *Pten-*deficient tumours) using the spatially restricted *ey-GAL4* driver. This model generates reproducible neoplastic growth while allowing animals to survive to adulthood, thereby providing an opportunity to investigate how localised tumours communicate with distant host tissues and ultimately influence disease outcome. Although tumour growth is confined to the dispensable eye primordium, *Pten*-deficient tumours caused substantial host lethality (∼40-60%). Moreover, whereas all larval eye discs displayed neoplastic overgrowth, only a subset of surviving adults developed overt eye tumours^18^.

Untargeted UHPLC–MS metabolomics identified tryptophan metabolism (Fig. 1a) as one of the most profoundly altered pathways in tumour-bearing larvae. Reanalysis of the kynurenine pathway revealed a striking, more than 200-fold increase of 3-hydroxykynurenine (3-HK; Fig. 1b), accompanied by systemic depletion of tryptophan (Fig. 1c). In contrast, kynurenic acid (KynA), which represents the alternative branch of the pathway (Fig. 1a), remained unchanged (Fig. 1b). These findings indicate that *Pten*-deficient tumours selectively redirect systemic tryptophan metabolism towards the 3-HK-producing branch rather than causing a general increase in kynurenine pathway activity.

Elevated kynurenine metabolites in tumour-bearing larvae originate primarily from the liver-like fat body, not the tumour itself: Expression of *tryptophan 2,3-dioxygenase* (*TDO/vermilion*)^25^ and *kynurenine 3-monooxygenase* (*KMO/cinnabar*)^26^ was strongly upregulated in the fat body, with only modest induction in tumours (Fig. 1d). Targeted GC–MS profiling confirms substantial 3-HK accumulation in the fat body (Fig. 1e), establishing it as the principal source of tumour-induced systemic kynurenine.

*Pten* knockdown alone, but not *Dl* overexpression alone, induced this metabolic signature (Fig. 1f), demonstrating that kynurenine pathway activation is a direct consequence of *Pten* loss and not merely secondary to tumour burden or developmental tissue disruption.

### Tumour-derived nitric oxide remotely activates tryptophan catabolism genes

Inflammation is a major inducer of the kynurenine pathway IDO and TDO enzymes^14^, while chronic inflammation via aberrant Nitric oxide (NO), a conserved regulator of innate immunity and inflammatory signalling^27^, is a hallmark of oncogenic PTEN loss in Drosophila cancer and human paediatric cells^18,21^. Intriguingly, increased *Nitric oxide synthetase* (*NOS*) is linked to increased *IDO2* expression in pancreatic cancer^28^. As, loss of *PTEN,* which causes constitutive PI3K–Akt pathway activation, directly upregulated Nos, leading to aberrant NO production, we explored whether Nos from tumours may be responsible for the transcriptional activation of tryptophan metabolism genes.

Consistently with increased tumour NO production, metabolomic data show that tumour-bearing larvae have reduced L-arginine, the substrate for NO synthesis (Fig. 2a,b), together with increased expression of the cationic amino acid transporter *Slimfast* (*Slif*) (Fig. 2c), which mediates arginine uptake^29^. Moreover, tumour-specific knockdown of *Slif* via RNAi significantly reduced tumour incidence, further supporting a role for amino-acid-dependent NO production in tumour progression^20^ (Fig. 2d). We next tested whether NO tumour-derived NO provides a signal linking oncogenic *Pten* loss to the systemic activation of tryptophan metabolism.

**Fig. 2.**
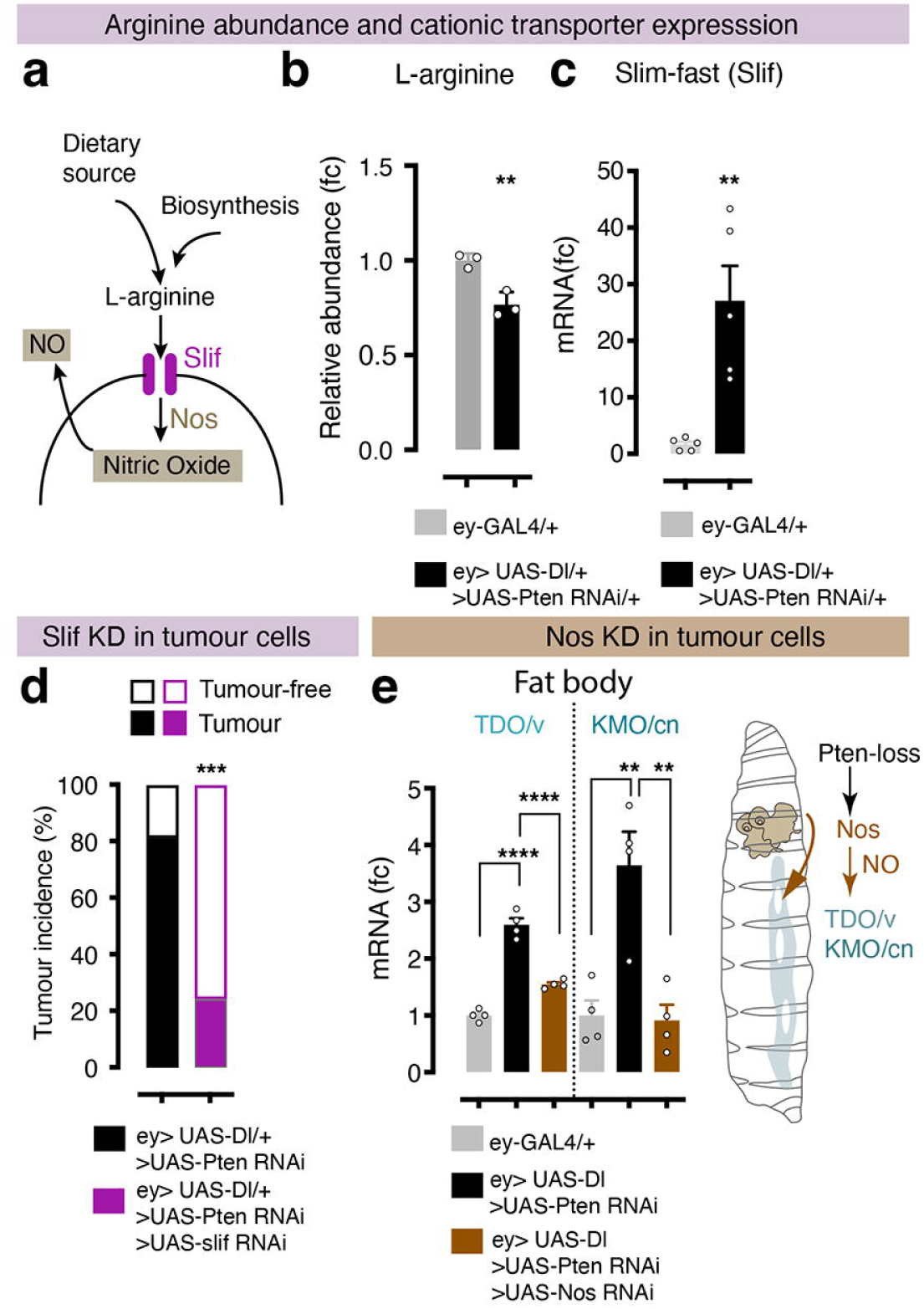
Tumour-derived nitric oxide remotely activates host kynurenine pathway TDO/v and KMO/cn genes. **a,** Scheme of the production of Nitric oxide from L-arginine. **b,** Metabolomic data show reduced global L-arginine in tumour larvae compared to control. Relative abundance (fold change to genetic control) Unpaired two-tailed Student t-test: *ey-GAL4* vs. *ey>Dl>Pten RNAi* (*n* = 3) *P =* 0.0060 (**). **c,** Expression of arginine transporter *Slif* gene was quantified in whole-larvae. Data are presented as relative expression (fold change vs control), with bars and errors showing mean ± s.e.m. and dots representing individual biological replicates. Statistical significance was calculated using an unpaired Student t-test: *P =* 0.0035 (**). **d,** Tumour-specific RNAi knockdown using *ey-GAL4* of the cationic amino acid transporter Slif reduced tumour incidence (*n* = 20 per genotype). Data are presented as mean ± s.e.m. Statistical was calculated using unpaired Student t-test: *P* = 0.0001 (***). **e,** Tumour-specific knockdown (KD) of *Nos* (*ey>Dl>Pten RNAi>Nos* RNAi) reduced expression of *TDO/v* and *KMO/cn* in the fat body. Expression was quantified in dissected fat bodies. Data show mean ± s.e.m. of fold change (fc) to control, with dots representing individual biological replicates (*n* = 4). Statistical significance was calculated using an ordinary one-way ANOVA with Tukey’s multiple comparison test with exact P values (left to right): *P* < 0.0001 (****), *P* < 0.0001 (****), *P* = 0.0033 (**), and *P* = 0.0027 (**).

Tumour-specific *Nos* depletion significantly reduced expression of not only the IDO-related enzyme gene, *TDO/v* but also the downstream enzymatic step, *KMO/cn* in the fat body cells (Fig. 2e), indicating that tumour-derived NO is required for host kynurenine pathway genes. These observations identify tumour-derived NO as a signal that remotely activates host tryptophan metabolic genes, providing a mechanistic link between oncogenic *Pten* loss and systemic production of the kynurenine metabolite 3-HK.

### Host-derived 3-HK promotes tumour progression

To determine whether systemic activation of the kynurenine pathway contributes to tumour progression, we inhibited 3-HK production genetically and pharmacologically. Tumour-specific knockdown of *TDO/v* or *KMO/cn* via RNAi modestly reduced tumour incidence, whereas systemic reduction, via endogenous mutations, strongly suppressed tumour growth (Fig. 3a), implicating host-derived kynurenine metabolism in promoting tumour progression.

**Fig. 3.**
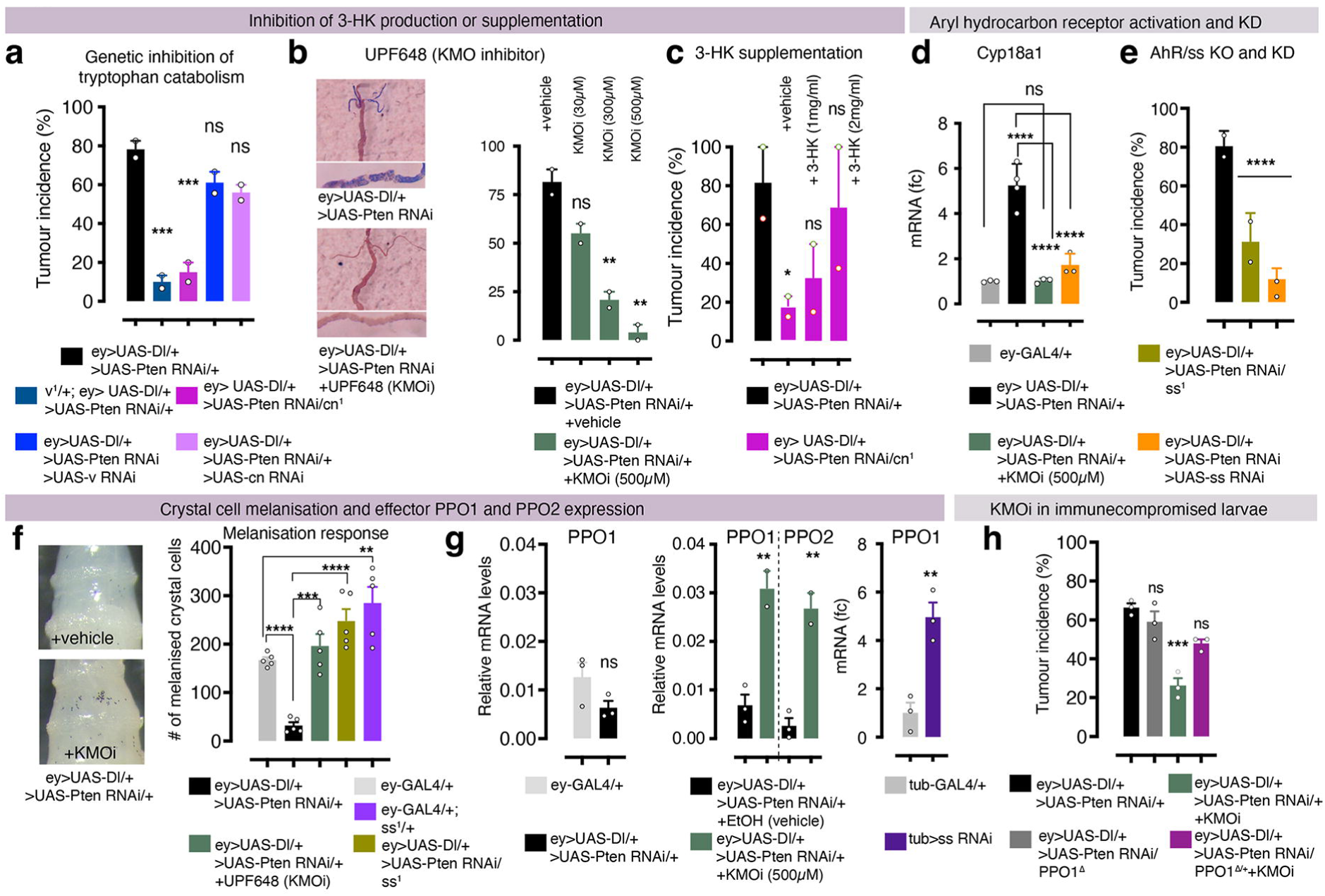
Host-derived 3-HK and AhR signalling promote tumour growth and suppress crystal cell immune function. **a,** Tumour incidence in adults derived from larvae with systemic inhibition (genetic mutation) or tumour-specific knockdown (KD) of TDO/*vermilion (v)* or KMO/*cinnabar (cn)* via RNAi expression. Bars show percentage of eye with tumours in the indicated genotypes, pooled from two independent crosses. As tumour incidence can vary across experiments, each biological repeat is normalised to their respective control run in parallel. *ey>Dl>Pten RNAi/+* (*n* = 46 eye scored), *v^1/+^* BDSC:137; *ey>Dl> Pten RNAi/+* (*n =* 30), *ey>Dl> Pten RNAi/+, cn^1^* BDSC:263/+ (*n* = 20), *ey>Dl> Pten RNAi/+>v RNAi* BDSC:50641 (*n* = 36), *ey>Dl> Pten RNAi/+>cn RNAi* BDSC:65029 (*n* = 50). For multiple comparisons, the ordinary one-way ANOVA with Tukey’s post hoc test was used. Exact *P*-values (left to right): *P* = 0.007 (***), *P* = 0.004 (***). **b,** Pharmacological inhibition of 3-HK synthesis with the KMO inhibitor UPF648 (or vehicle, EtOH). Left: Malpighian tubules from female *ey>Dl>Pten RNAi* larvae, showing that excess 3-HK are stored in this organ leading to pigmented tubules—an effect completely abolished by KMO inhibitor treatment. Bar plots show the percentage of eye tumour, pooled from two biological replicates. Statistical analysis using Ordinary one-way ANOVA with Tukey’s multiple comparison test (left to right): *P =* 0.0036 (**), *P* =0.0014 (**). ey*>Dl>Pten RNAi* + EtOH vehicle (*n* = 26)*; +* KMOi 30 µM (*n* = 36); *+* KMOi 300 µM (*n* = 28); *+* KMOi 500 µM (*n* = 32). **c,** Dietary supplementation with 3-HK in tumour larvae with genetic impairment of KMO*/cn* (*cn^1^*/+). Larvae were reared on food supplemented with increasing concentrations of the metabolite 3-HK or vehicle (H_2_O), and tumour incidence was scored in adults and compared with animals without *cn^1^* mutation. Bars show percentage of eyes with tumour, pooled from two experiments. *ey>Dl>Pten RNAi/+* (*n* = 60), *ey>Dl> Pten RNAi/+, cn^1^* BDSC:263/+ vehicle (*n =* 42), *ey>Dl> Pten RNAi/+, cn^1^* BDSC:263/+ 3-HK (1 mg/mL; *n =* 40), *ey>Dl> Pten RNAi/+, cn^1^* BDSC:263/+ 3-HK (2 mg/mL; *n =* 26). Ordinary one-way ANOVA with Tukey’s post hoc test was used. *P* = 0.0212 (*). **d,** Expression of the AhR/spineless (ss) target gene *Cyp18a1* is elevated in tumour-bearing eye discs. This induction is reduced by KMO inhibition (500 µM UPF648) and by silencing AhR/*Spineless* via RNAi. Data are presented as relative expression (fold change vs control), with bar plots showing mean ± s.e.m. and dots representing individual biological repeats (*n* = 3). Statistical significance was assessed using an ordinary one-way ANOVA with Tukey’s multiple comparison test. *P <* 0.0001 (****). **e,** Tumour incidence rescue by reducing ss gene copy (*ss^1^*, BDSC:2973) or tumour-specific inhibition of AhR/Spineless (ss) using RNAi (*UAS-ss RNA-i*, v10832). Ordinary one-way ANOVA with Forsythe test: *P* < 0.0001 (****) (*n* = 30). **f,** Systemic crystal cell melanisation was quantified as the number of melanised foci per larva following heat induction (*n* = 5 larvae). Data show mean ± s.e.m. Unpaired Student *t-*test for pairwise comparisons (left to right): *P* < 0.0001 (\*\*\**\**), *P =* 0.0002 (\**\*\**), *P* < 0.0001 (****), and *P* = 0.0077 (**). **g,** Expression of crystal cell effector PPO1 or PPO2 in the indicated genotypes. Data show mean s.e.m., and dots represent biological repeats. Relative mRNA expression is shown for comparison of *PPO1* and *PPO2*, or normalised to the control (fold change, fc) for *PPO1* alone. Pairwise comparisons used unpaired *t-test* (left to right): *P =* 0.0089 (**), *P =* 0.0046 (**). **h,** Graph shows tumour incidence in animals treated during larval development with KMO inhibitor (500 µM) and with partial genetic inactivation of *PPO1* via mutation *(PPO1*^Δ^/+), demonstrating that melanisation contributes functionally to the anti-tumour effect. Data represent the percentage of flies with tumour in the indicated genotypes and conditions. Ordinary one-way ANOVA, Dunnett’s multiple comparisons test: *P* = 0.0004 (***).

Pharmacological inhibition of KMO with the selective inhibitor UPF648^30^ markedly reduced Malpighian tubule pigmentation, a surrogate for systemic 3-HK accumulation^31^, confirming effective inhibition of the pathway (Fig. 3b). KMO inhibition also suppressed tumour formation in a dose-dependent manner (Fig. 3b), further supporting a tumour-promoting role for host-derived 3-HK.

To establish that 3-HK is the relevant downstream metabolite, we supplemented the diet with exogenous 3-HK. Dietary 3-HK restored tumour growth in *KMO/cn¹* heterozygotes in a dose-dependent manner (Fig. 3c). Together, these findings identify 3-HK as the principal tumour-promoting kynurenine metabolite.

Kynurenine metabolites can signal via the aryl hydrocarbon receptor (AhR), a ligand-activated transcription factor integral to detoxification, host defence, and immune tolerance across vertebrates^15^. We therefore examined the *Drosophila* AhR orthologue *spineless* (*ss*)^34,35^. Expression of the canonical AhR/Spineless target gene *Cyp18a1* was upregulated in the tumour discs and reduced following either KMO inhibition or AhR/Spineless silencing (Fig. 3d), indicating tumour-induced activation of AhR/Spineless signalling. Consistent with this model, systemic and tumour-specific knockdown of *spineless* reduced tumour growth (Fig. 3e), placing AhR downstream of 3-HK, indicating that AhR signalling is required downstream of 3-HK to promote tumour progression.

### KMO inhibition suppresses tumour growth by restoring crystal cell effector activity

*Drosophila* crystal cells mediate a major cytotoxic arm of insect innate immunity through PPO1 and PPO2, generating melanin, reactive quinones and reactive oxygen species that eliminate pathogens and abnormal cells but can also damage host tissues if not tightly regulated^23,37^. Whereas hyperplastic *ey>Dl* tissues robustly activated this cytotoxic programme, *Pten*-deficient tumours failed to do^18^ despite the later representing a more severe neoplastic challenge. We therefore asked whether kynurenine signalling promotes tumour progression by suppressing crystal cell effector activity.

Directly linking kynurenine metabolism to innate immunity, pharmacological inhibition of KMO/cn restored crystal cell melanisation in tumour-bearing larvae (Fig. 3f) and reactivated expression of the crystal cell effectors *PPO1* and *PPO2* (Fig. 3g). Likewise, inhibition of AhR/Spineless signalling restored melanisation (Fig. 3f) and reducing AhR/Spineless in healthy animals was sufficient to increase crystal cell effector *PPO1* expression (Fig. 3g). These findings identify the NO–KMO (3-HK)–AhR pathway as an active suppressor of crystal cell cytotoxicity, analogous to the immunotolerogenic role of kynurenine signalling in mammals^38^.

To functionally demonstrate that restoration of crystal cell activity contributes to the tumour-suppressive effect of KMO inhibition, we reduced crystal cell effector function using the *PPO1*^Δ^ null mutation. *PPO2* was not examined genetically because, in addition to its role in melanisation^37^, it is required for systemic oxygen transport and normal growth^39^, which would confound tumour analysis. Notably, the partial loss of *PPO1* significantly attenuated the tumour-suppressive effect of KMO inhibition (Fig. 3h), demonstrating that reactivation of crystal cell effector activity is an essential component of KMO inhibitor-mediated tumour suppression.

Collectively, these findings support a model in which tumour-induced NO–KMO (3-HK)–AhR pathway promote an unresponsive or tolerogenic haemocyte state, characterised by suppressed terminal effector functions. The cellular basis of this immune remodelling is examined below.

### Oncogenic *Pten*-deficient tumours remodel the immune microenvironment towards immune suppression

To determine how tumour-derived NO and host 3-HK are associated with remodelling of the tumour immune microenvironment, we characterised tumour-associated haemocyte populations and compared them with those associated with healthy and hyperplastic tissues.

Resident haemocytes were abundant in control eye–antennal discs. These populations consisted predominantly of plasmatocytes (macrophage-like cells), together with approximately 5% crystal cells (Fig. 4a,b) and a small population of lamellocytes (∼2.7–3%), broadly resembling the proportions reported for circulating and sessile haemocytes. Upon encountering abnormal or damaged cells, crystal-cell undergo rupture to release cytotoxic material during killing of abnormal or damaged cells^23,37^ (Fig. 4c,d). Terminal differentiated plasmatocyte expressing NimC1, a phagocytic receptor^40^, were frequently associated with crystal-cell debris (Fig. 4e), consistent with local immune surveillance during tissue homeostasis.

**Fig. 4.**
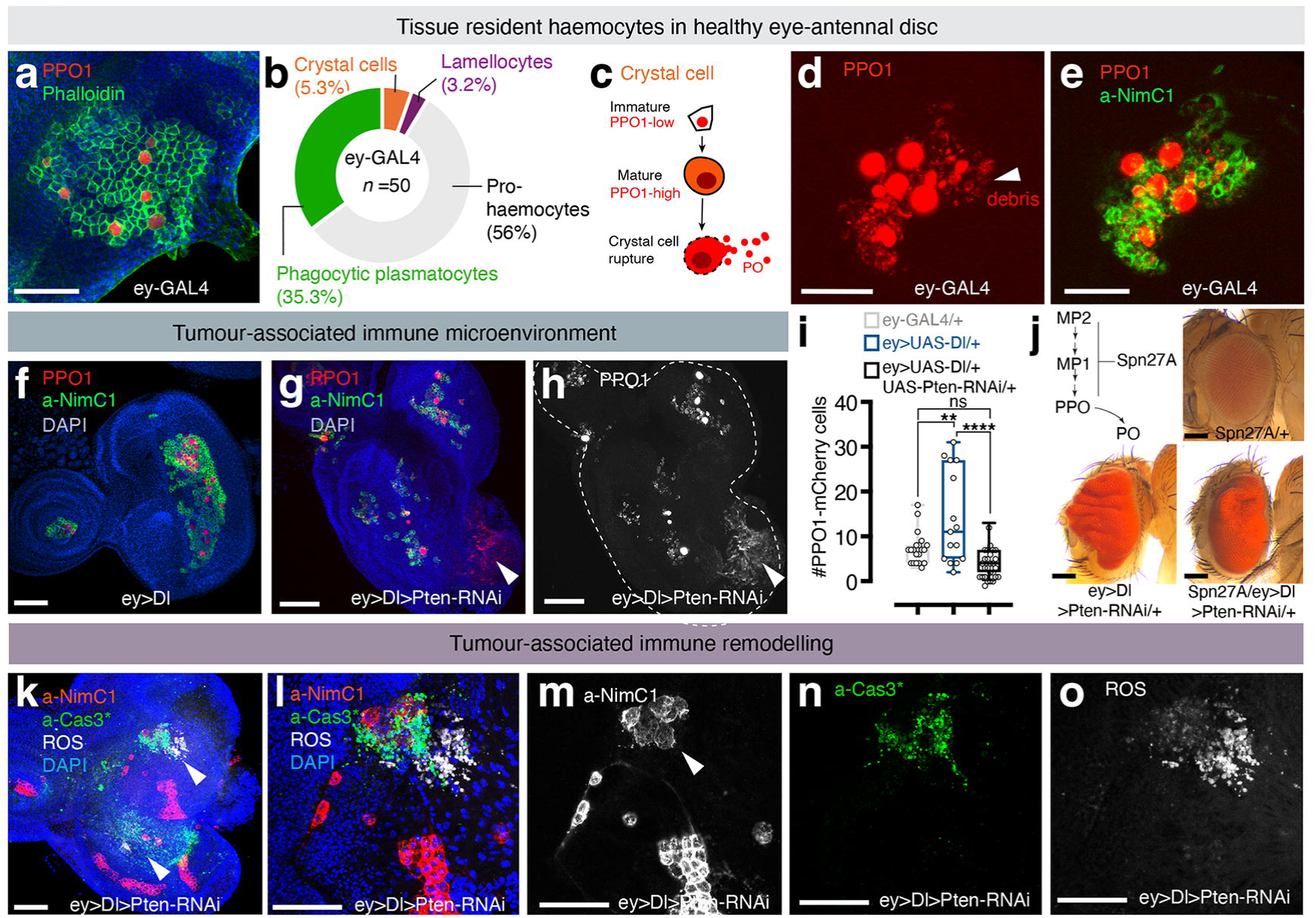
Tissue resident immune populations and the tumour-associated immune remodelling. **a, d–h k–n,** Confocal images of resident haemocyte populations in control (**a**, **d, e**), hyperplastic (**f**), and tumour (**g, h, k–o**) eye-antennal discs. All discs are from female larvae. In **a,** resident haemocytes are identified by F-actin (Phalloidin, green), crystal cells by PPO1-mCherry^55^ (also known as Bc-F6-mCherry; red) and nuclei are counterstained with DAPI (blue). **b**, Quantification of resident haemocyte subtypes in control discs (pooled from *n =* 50 discs). **c**, Schematic of crystal cell maturation leading to cellular rupture^51^. **d,** Intact crystal cells (PPO1-mCherry) and mCherry+ debris in a representative control disc. Debris visualised by image saturation. **e**, Phagocytic plasmatocytes (NimC1+, green) aggregate around mCherry+ debris. **f,** Hyperplastic *ey>Dl* eye-antennal discs show large aggregated of NimC1-positive plasmatocytes (green) and PPO1-mCherry (red). Nuclei are counterstained with DAPI (blue). **g**,**h**, Confocal image of a representative tumour discs, showing tumour-associated crystal cell (PPO1-mCherry: red in **g** and grey in **h**) and NimC1-positive plasmatocytes (green). Nuclei are counterstained with DAPI (blue). **h,** Image is saturated to illustrate mCherry debris and a low PPO1-positive area devoid of plasmatocytes (arrowheads in **g** and **h**). **i,** Quantification of crystal cells per disc by genotype: control *ey-GAL4/+* (*n* = 20), *ey>Dl/+* (*n* = 15), *ey>Dl>Pten RNAi/+* (*n* **=** 27). Box-and-whiskers plots indicate the median, interquartile range, and the minimum and maximum values, with individual values as dots. Ordinary one-way ANOVA, Tukey’s multiple comparison tests: *P* = 0.0041 (**), *P* < 0.0001 (****). **j,** Scheme: The Serpin 27A immune checkpoint prevents excessive activation of the melanisation cascade involving a proteolytic pathway converting PPO1 and PPO2 to active phenoloxidases (PO1 and PO2). Representative adult eyes of the indicated genotypes. **k,** Whole-tumour view and (**l–o**) magnified and single-staining images for ROS (grey in **k** and **l**), cleaved caspase3 (green in **k,l,n**), and NimC1 (red: **k, l,** and grey: **m**) Arrowheads point to high ROS labelling in **k** and the associated enlarged plasmatocytes in **m**. Discs are outlined by white lines in **c,-d,f-i,n,o**. Scale bars, 50 µm, except 20 µm in **a, d-e** and 100 µm in **j.**

To establish a benchmark anti-tumour immunity, we examined benign eye-disc hyperplasia (*Dl* overexpression alone) and observed that there was a robust immune response: a marked accumulation of crystal cells and large aggregates of activated NimC1-positive plasmatocytes (Fig. 4f), in agreement with increased expression of crystal cell effectors *PPO1* and *PPO2* assessed by qPCR^18^. Genetic reduction of *PPO1-PPO2* activity converts this overgrowth to neoplasia^18^, underscoring the importance of crystal cells in restraining preneoplastic tissue expansion.

In stark contrast, oncogenic *Pten*-deficient tumours elicit an attenuated crystal cell– plasmatocyte response (Fig. 4g,h). Crystal-cell numbers per tumour are similar to healthy controls, but a much lower than in hyperplastic *ey>Dl* discs (Fig. 4i). Suppressed terminal immune effector activity is evident, with areas of low PPO1-mCherry-positive cells and absence of plasmatocytes (Fig. 4g,h). Reducing *Serpin 27A* dosage, an immune checkpoint of melanisation^41^, decreases tumour burden (Fig. 4j), reinforcing the significance of the PPO-mediated anti-tumour response for control of tissue expansion.

Tumour-associated NimC1-positive plasmatocytes progressively lost their characteristic clustered organisation, accompanied by the emergence of isolated enlarged cells with reduced NimC1 expression (Fig. 4k-o and see later, Fig. 5a), consistent with immune remodelling states. Enlarged NimC1-positive plasmatocytes were associated with bursts of ROS and cleaved caspase-positive cells (Fig. 4l–o), indicating local oxidative stress and tissue damage in the vicinity of these remodelled haemocytes.

**Fig. 5.**
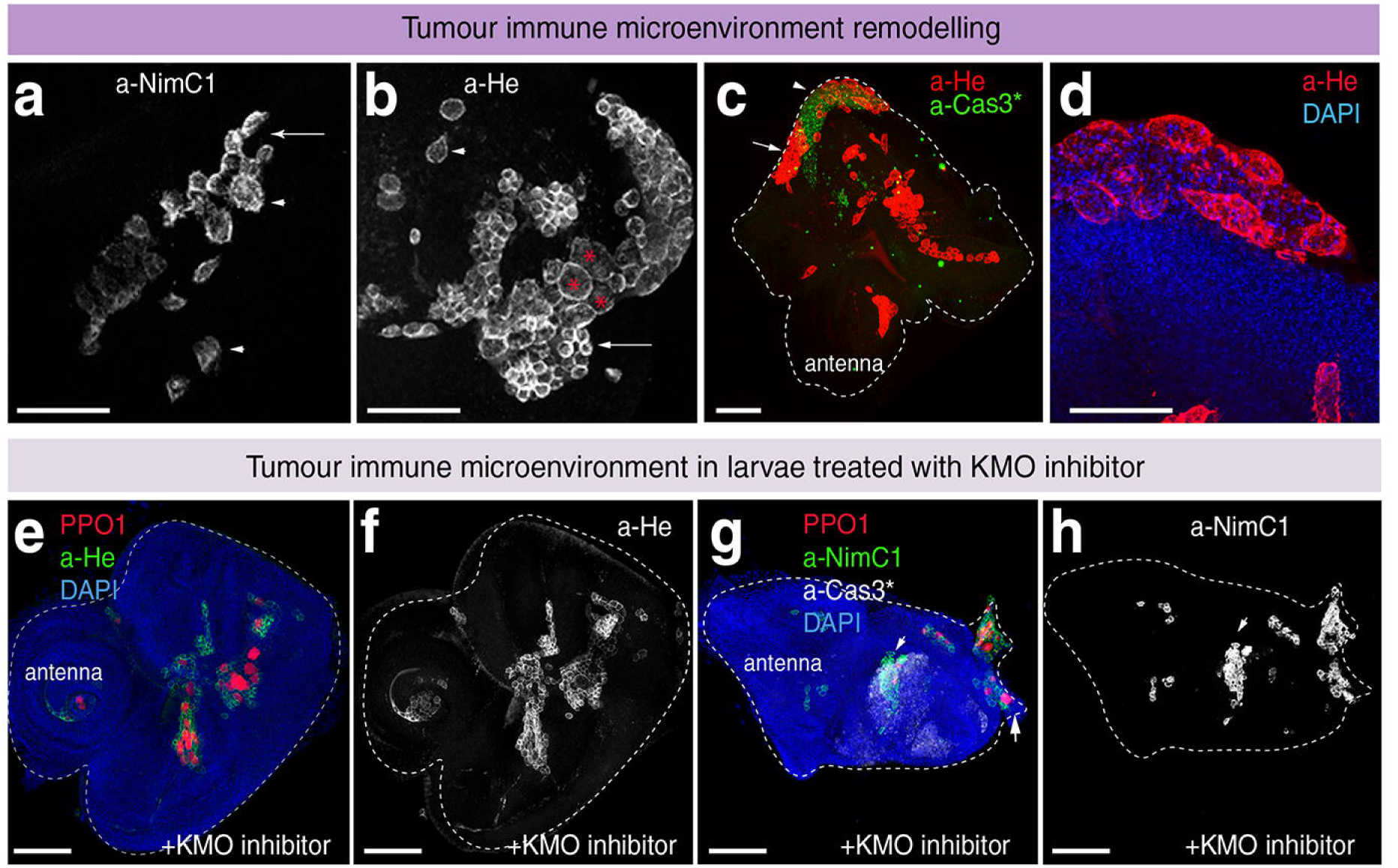
Tumour-induced immune remodelling is reversed by KMO inhibition. All images show tumour eye–antennal discs (*ey>Dl>Pten RNAi/+*); genotypes are omitted from the images for clarity. **a,** High-magnification image showing remodelling of NimC1-positive plasmatocytes within the tumour microenvironment (*n* = 15 discs). **b,** High-magnification image showing haemocyte remodelling, with giant haemocytes (asterisks) labelled by anti-Hemese (a-He) emerging from aggregates of small haemocytes (arrow, *n* = 20). **c,** Whole-tumour views showing giant haemocytes localised to regions with extensive cleaved-caspase staining (arrowhead), whereas aggregates of small haemocytes are largely devoid of caspase staining (arrow, *n* = 15). **d,** Higher magnification of giant haemocytes in **c**, revealing internal membrane structures and multiple DAPI-positive puncta. **e–h,** Tumour discs following KMO inhibitor treatment (*n* = 15), showing crystal cells (PPO1-mCherry; red in **e,g**), pan-haemocytes (a-He; green in **e** and grey in **f**) and phagocytic plasmatocytes (NimC1; green in **g** and grey in **h**). **f,** Single-channel a-He image illustrating restoration of small-haemocyte aggregates. Following KMO inhibition, cleaved-caspase staining (grey in **g**) is detected within small-plasmatocyte aggregates (arrow). **h,** Single-channel image of NimC1-positive plasmatocytes. **c,e-h** Scale bars, 50 µm, except 20 µm in **a, b,e.**

As NimC1 staining declines (Fig. 5a), we used the pan-haemocyte marker Hemese^42^ revealing a striking transition towards giant haemocytes (Fig. 5b) that segregated from aggregates of conventional plasmatocytes and accumulated at sites of chronic tissue damage (Fig. 5c-d). These giant haemocytes reached 40–60 μm in diameter and did not express NimC1. These exhibit stinking features such as they frequently contained extensive internal membranes and fragmented or multiple DAPI-labelling foci nuclei (Fig. 5e), reminiscent to tumour -associated multinucleated giant macrophages in mammal^43^.

Importantly, KMO inhibition completely eliminated or reduced the abundance of the emergence of altered giant haemocytes (Fig. 5f,g), while restoring immune activity (Fig. 5h,i), supporting key roles for 3-HK in immune compartment remodelling.

## Discussion

Our study identifies a tumour–host metabolic circuit linking *Pten* loss to systemic kynurenine metabolism, driving immune tolerance and tumour progression. Tumour-derived nitric oxide induces host 3-hydroxykynurenine production, which, together with AhR signalling, shapes an immunosuppressive environment. This restrains the scalation of cytotoxic melanisation effector functions — limiting immune-mediated collateral tissue damage while promoting tumour growth. Immune tolerance thus emerges as a dynamic, host-wide metabolic adaptation that coexists with basal immune surveillance.

Comparisons between tumours, preneoplastic hyperplasia, and controls reveal a shift towards immune suppression during tumour progression, accompanied by reduced immune-cell abundance and extensive remodelling of myeloid-like immune populations. Current models propose that tumour-derived damage signals, including ROS and MMP1, recruit circulating plasmatocytes, which can promote or suppress tumour growth or have little measurable effect^5,44–47^. We identify a substantial population of resident haemocytes in healthy eye tissue, establishing a local immune-surveillance compartment before tumour initiation. These cells remain plastic and diversify in response to neoplastic transformation. Tumour progression therefore involves both remodelling of resident haemocytes and recruitment of circulating cells. The spatially organised *Drosophila* eye tumour model provides an opportunity to resolve these dynamics and, in particular, the emergence of spatially compartmentalised tumour–immune microenvironments.

These findings have broad implications because *PTEN* loss is frequent in human cancers and in cancer predisposition syndromes such as Cowden disease^53^, and is associated with failure of immune tolerance^19^, suggesting that *PTEN* loss in human cancer may activate alternative immune checkpoints via tryptophan metabolism. The genetic amenability of *Drosophila Pten*-loss models can provide a powerful, spatially resolved platform to dissect further how tryptophan metabolic branches regulate tumour–immune–metabolic interactions and shape host immunometabolism and segregated tumour microenvironments.

Together, our findings uncover a tumour–host metabolic–immune circuit in which oncogenic *Pten* loss remotely reprogrammes host tryptophan metabolism to promote immune tolerance and tumour progression. Importantly, this systemic remodelling is already detectable following *Pten* loss in the absence of an overt tumour, providing evidence that specific oncogenic lesions can precondition the host metabolic and immune environment before tumour establishment. Thus, tumour–immune diversity may arise not only through progressive adaptation to an evolving tumour microenvironment, but also from systemic states preconfigured by the initiating oncogenic lesion. By identifying KMO-dependent 3-HK production as an actionable node connecting tumour genotype, systemic metabolism and innate immunity, our study highlights host metabolism as a determinant of tumour outcome. It suggests that targeting these early tumour–host circuits may provide opportunities to prevent or reverse innate immune tolerance.

## Methods

### Fly strains and husbandry

*Drosophila melanogaster* were maintained on standard Iberian medium^56^ (at 25 °C under a 12 h light/12 h dark cycle unless otherwise indicated. Experimental crosses were performed at 26.5 °C. The following strains were used: Oregon-R, Canton S, *w^1118^*, *ey-GAL44/CyO-twist-GFP*, *ey-GAL4>UAS-Dl/CyO-twist-GFP*, *ey-GAL44/CyO-tub-GAL80, UAS-Dl::mRFP/TM6B* (BSDC:26696), *UAS-Pten RNAi* (BSDC: 25967, P{TRiP.JF01987}attP2),, *UAS-Pten RNAi* (BDSC:8550, BDSC:8548), *UAS-Nos RNAi* (BDSC:50675, TRiP.HMC03076), *UAS-v RNAi* (B50641, TRiP.HMC03041 and v107798, P{KK108195}VIE-260B), *UAS-cn RNAi* (BDSC:65029; TRiP.HMC05903 and v105854 KK101938), *UAS-ss RNAi* (VDRC: 108732, KK107561), *UAS-slif RNAi* (BDSC:64972, TRIP.HMC05846), *UAS-mCD8-GFP* (BDSC:5137)-Additional mutant alleles used: *v^1^* (BDSC:137), *v^48a^* (BDSC: 4447), *v^36F^*(BDSC:142) *cn^1^* (BDSC:263), *cn^35k^* (BDSC:268), *ss^1^* (BDSC:2973),, *PPO1*^Δ^ (BDSC:56204); *spn27A^ex32^/CyO Act-GFP* (BDSC:67098), *PPO1-mCherry.F6* (BDSC: 600219).

Unless otherwise specified, experimental crosses were performed with two to three independent biological replicates. Because the *vermilion (v)* and *white* loci are widely used as selectable markers in *Drosophila* transgenesis, key findings were validated using complementary genetic and pharmacological approaches (see Extended Data Fig. 1a). The effects of *v* knockdown were independently confirmed using mutant alleles, with hypomorphic alleles producing less strong rescue: *v^1^* (null: 8.8% tumour incidence, *v^48a^*: 48%, and *v^36f^*: 62.28 %, tumour “control”: 80%). The effect of *cinnabar (cn)* knockdown (*cn^1^*) was reproduced pharmacologically using the KMO inhibitor UPF648 and reversed by dietary supplementation with 3-hydroxykynurenine (3-HK). Data are shown for null alleles. The tumour-promoting effect of *white* knockdown (RNAi) was independently reproduced by knockdown of the related ABC transporter *scarlet*. Tumour phenotypes were consistent across the independent tumour models used in this study as presented previously^18^. Increasing copies of UAS-transgenes did not diluted the tumour phenotype as verified including a *UAS-GFP-RNAi*. The effect of *UAS-Pten RNAi* transgenes generated in both *y v (TRiP)* and *w* backgrounds on tumorigenesis was consistent and recapitulated by Akt overexpression^18^. Because the second chromosome balancers *CyO* and *SM6b* carry the weak *cn²* allele, progeny inheriting these balancers were excluded from tumour incidence and tumour burden analyses.

### Tumour model and tumour incidence

Oncogenic *Pten*-deficient eye tumours were generated by driving *UAS-Delta* (*UAS-Dl* or *UAS-Dl::RFP*, for some larval analyses) and *UAS-Pten RNAi* with eyeless (*ey)-GAL4*, as previously described^18,21^. This combination induces neoplastic growth in the eye imaginal disc through synergistic activation of the Notch and PI3K–Akt pathways. Tumour incidence was determined in adult survivors by stereomicroscopic examination of eye morphology. Flies displaying a visible eye tumour were scored as tumour-positive, and tumour incidence is reported as the percentage of tumour-positive adults among all surviving adults bearing the oncogenes. Because tumour penetrance varies between experimental cohorts, intervention effects are presented relative to the corresponding control group analysed in parallel.

### Untargeted metabolomics (UHPLC–QTOF–MS)

Whole L3 larvae at the wandering stages (*n* = 15 in three biological replicates per condition) were homogenised in methanol:chloroform:water (3:1:1, v/v/v) using 220mg of glass beads (QIAGEN, ref.#13116-400) and the TissueLyser-LT cell disruptor (QIAGEN, ref.# 85300). Extracts were centrifuged and stored at −20 °C. Samples were analysed on an Agilent 1290 Infinity UHPLC coupled to a 6540 UHD Q-TOF (positive ESI mode). The samples were injected randomly in order to eliminate any drift effects that the equipment or analysis conditions might present. Data were processed using MassHunter and Mass Profiler Professional. Features were filtered to remove signals <3× blanks and present in <66% of samples. Data were normalized to total ion signal and Pareto-scaled prior to multivariate analysis. PCA and statistical analyses were performed using MetaboAnalyst. Zero or missing variables were replaced with small values (the half of the minimum positive values in the original data) assuming that the missing values are caused by low abundance metabolites (i.e., below the detection limit) in the control animals at this stage. Metabolite annotation was based on accurate mass matching against METLIN and KEGG and is reported as putative unless validated with standards.

### Targeted metabolomics (GC–MS)

Whole larvae (15 wandering L3 per condition), haemolymph (extracted from 50 wandering L3 larvae), and fat bodies (from 30 larvae) were flash-frozen and extracted in 90% methanol containing succinic-d4 acid (internal standard)^57^. Samples were derivatized with methoxylamine hydrochloride and MSTFA. GC–MS analysis was performed on an Agilent 7890B Gas Chromatograph coupled to a 5977 MS (50–500 m/z) (Agilent Technologies). The mass spectrometer was set to execute a SIM/SCAN acquisition mode over a mass range of 50–500 m/z, allowing to the high sensitivity identification of the metabolites of interest. Quantification was normalised to internal standard and tissue mass.

### Tissue collection, RNA isolation and qRT–PCR

Wandering third-instar (L3) larvae were dissected in cold PBS in biological triplicate.. For other tissue-specific analyses, ten fat bodies or 25 eye imaginal discs were pooled per sample. For whole-larva RNA extraction, five larvae per sample were collected. Samples were preserved in RNAlater and stored at −80 °C. Total RNA was extracted using the RNeasy Mini Kit (Qiagen ref. # 74106) and treated with TURBO DNase (Applied Biosystems, ref. # AM1907). RNA quantity and purity were assessed by NanoDrop ND-1000 spectrophotometry. One microgram of total RNA was reverse-transcribed using SuperScript III (Invitrogen, ref.#58063) with oligo(dT) (Invitrogen, ref.#58063) and random hexamer primers (Invitrogen, ref.#100026484). Reactions were incubated at 50 °C for 60 min followed by heat inactivation. Quantitative PCR was performed using Power SYBR Green Master Mix (Applied Biosystems, ref.# 4367659), on a 7500 Real-Time PCR system Applied Biosystems. Cycling conditions were: 95 °C for 10 min; 40 cycles of 95 °C for 15 s and 60 °C for 40 s. Relative expression levels were calculated using the ΔΔCt method and normalised to *Rp49*. Each biological replicate was analysed in technical triplicate. Fold changes were expressed relative the internal control. Primer sequences are listed in Table 1.

### Dietary and Pharmacological supplementation

Dietary and Pharmacological supplements were added at the following final concentrations: 3-Hydroxykynurenine (Sigma-Aldrich, ref. #H1771): 1–2 mg/mL; ; KMO inhibitor (UPF648, ref. #4926/10, Tocris Bioscience, Bristol, UK): 30–500 µM; TDO inhibitor (680C91, ref. #4392) was used up to 100 µM and it caused lethality at all concentrations tested. Vehicle concentrations were kept constant across conditions.

### Immunostaining, image acquisition, and analysis of tissue- and tumour-associated haemocytes and stromal cells

Larval eye discs were fixed in 4% paraformaldehyde, permeabilised, and incubated with primary antibodies against cleaved Caspase 3 (CST ref#9661, rabbit anti-Caspase 3, 1:100). Distinct haemocyte classes were labelled using mouse anti-NimC1 (Dm N1/9, 1:50) and anti-Hemese/H2 (Dm 1.2/1, 1:50, labelling ∼80% of haemocytes^42^) from DSHB. Secondary antibodies (1:500), with or without phalloidin conjugates (Phalloidin Invitrogen-488 ref12379 1:100; Rhodamine Ref R415 1:500 and -647 A22287 1:100), were applied. Samples were mounted in Vectashield® antifade medium (Vectorlabs, H-1200-10) with DAPI and imaged under identical confocal settings. Eye disc area, fluorescence intensity, and haemocyte populations were quantified using ImageJ. To assess resident haemocyte precursors, Hemese, and F-actin was used. Plasmatocytes represent a molecularly diverse haemocyte class^59-61^. Matured phagocytic plasmatocytes were labelled by NimC1. All imaginal discs shown are from female larvae. Mean tumour size was calculated using ImageJ.

### Crystal cell melanisation assay

To test innate immune cell melanisation activity, wandering L3 larvae (112–120 h AEL) were heat-shocked at 60 °C for 10 min to induce melanisation reaction of active crystal cells. Images were acquired using a Leica M125 stereomicroscope. Crystal cells were counted in five larvae per genotype. In all of the images shown, the anterior is left, and the dorsal side is up.

### 3-Hydroxykynurenine accumulation

Kynurenine metabolites are stored or accumulate in the Malpighian tubules, which act as the secretory organs in insects. Tubules and guts were dissected from wandering L3 larvae (120– 140 h AEL) in Ringer’s solution. Female tubules were imaged, and inverted colour was applied in Adobe Photoshop (Fig. 3b).. Yellow pigmentation is also visible in tumour-bearing larvae through their transparent cuticle.

### ROS detection (CellROX)

Dissected eye disc from control and tumour larvae were incubated *ex vivo* in Scheider’s Drosophila medium (Gibco, ref #21729924) containing CellROX Deep Red Reagent kit (Life Technologies, ref #C10422) at 1:500 for 15 min in agitation at room, washed 3 times with PBS1x, fixed for 15min in 4% PFA, and mounted using Vectashield with DAPI, or followed by antibody staining as described above.

### Statistical analysis

Data are presented as mean ± s.e.m. unless otherwise indicated. Continuous variables were analysed using two-tailed Student’s t-test or one-way ANOVA with post hoc correction. Categorical data (tumour incidence, viability) were analysed using chi-square or Fisher’s exact test as appropriate. Statistical analyses were performed using R, GraphPad Prism 10.8 software (GraphPad Software), and MetaboAnalyst. P < 0.05 was considered statistically significant. Outliers were excluded only when predefined technical criteria were met. No statistical methods were used to pre-determine sample sizes, but our sample sizes are similar to those reported in previous publications^18^. The exact P values and sample size (n) are provided in the Figure legends. Figures created in Adobe Illustrator 2026 (version 30.6).

## Data availability

Metabolomics data are available via MetaboLights^62^ with identifiers MTBLS15287 and MTBLS15286. Reagents, *Drosophila* strains, custom scripts, and detailed protocols are available from the corresponding author upon reasonable request.

## Code availability

No new code was generated during this study.

## Acknowledgements

We thank the Bloomington Drosophila Stock Center (NIH P40OD018537), the TRiP consortium at Harvard Medical School (NIH/NIGMS, the Vienna Drosophila Resource Centre (VDRC, for stocks used in this study, and the Developmental Studies Hybridoma Bank at the University of Iowa for antibodies. Some Figures drawings were created with BioRender.com. We thank F. Serra, T. Xie, E. Sucena, M. Crozier, I. Aldo, S. Bray, for strains and reagents. We thank Carlos Canseco for technical support with LC-MS metabolomics, and the Genomics Unit at the CRG for assistance with the sequencing, M. Diaz for graphical illustrations, and L. Mira for assistance in fly stock keeping and behavioural assays. J.M.T. is supported by the National Institute of General Medical Sciences of the National Institutes of Health under a R35 Maximizing Investigators’ Research Award (MIRA; 1R35GM119557). The Spanish National Grants (<u>PID2019-106002RB-I00</u>) co-financed by the ERDF, the Proof of concept (PDC2022-133387-I00), the Asociación Española Contra el Cáncer (AECC) (<u>CICPF16001DOMÍ</u>), and Generalitat Valenciana Grant (PROMETEO/2021/027) co-financed with ERDF fund to M.D. M.D. was also funded by the “Severo Ochoa” Program for Centers of Excellence in R&D (CEX2021-001165-S) co-financed by ERDF. E. S-C. was a Spanish doctoral FPI fellow (PRE2018-085912) from the Spanish Ministerio de Economia y Competitividad and ML.U was supported by The Fundación General CSIC-ComFuturo programme, which has received funding from the European Union’s Horizon 2020 research and innovation programme under Marie Sklodowska-Curie gran agreement No. 101034263.

## Author contributions

Conceptualization: L.G-L, E. S-C, and M.D. Investigation: L.G-L, E.S-C, ML.U and M.D. Methodology: L.G-L, E. S-C, ML.U, C.G-E, E.B-I, JM.T and M.D. Writing—original draft preparation: L.G-L and M.D. Writing—review and editing: E.S-C, ML.U, JM.T and M.D. Supervision: L.G-L, E.S-C, and M.D co-supervised L.G-L and JM.T., Funding acquisition: ML.U, JM.T and M.D.

## Correspondence and requests for materials should be addressed to

Maria Dominguez.

